# Single Prolonged Stress Reduces Intrinsic Excitability and Excitatory Synaptic Drive onto Pyramidal Neurons in the Infralimbic Prefrontal Cortex of Adult Rats

**DOI:** 10.1101/2021.03.16.435686

**Authors:** Nawshaba Nawreen, Mark L Baccei, James P Herman

**Affiliations:** Dept. of Pharmacology & Systems Physiology, University of Cincinnati, Cincinnati, Ohio 45237-0506, United States; Neuroscience Graduate Program, University of Cincinnati, Cincinnati, Ohio 45237-0506, United States; Department of Anesthesiology, Pain Research Center, University of Cincinnati Medical Center, Cincinnati, Ohio 45237-0506, United States; Veterans Affairs Medical Center, Cincinnati, Ohio 45221, United States

**Keywords:** Prefrontal Cortex, Excitability, Stress, GABA, Glutamate

## Abstract

Post-traumatic stress disorder (PTSD) is a chronic, debilitating mental illness marked by abnormal fear responses and deficits in extinction of fear memories. The pathophysiology of PTSD is linked to decreased activation of the ventromedial prefrontal cortex (vmPFC). This study aims to investigate underlying functional changes in synaptic drive and intrinsic excitability of pyramidal neurons in the rodent homolog of the vmPFC, the infralimbic cortex (IL), following exposure to single prolonged stress (SPS), a paradigm that mimics core symptoms of PTSD in rats. Rats were exposed to SPS and allowed one week of recovery following which brain slices containing the PFC were prepared for whole-cell patch clamp recordings from layer V pyramidal neurons in the IL. Our results indicate that SPS reduces spontaneous excitatory synaptic drive to pyramidal neurons. In addition, SPS decreases the intrinsic membrane excitability of IL PFC pyramidal cells, as indicated by an increase in rheobase, decrease in input resistance, hyperpolarization of resting membrane potential, and a reduction in repetitive firing rate. Our results suggest that SPS causes a lasting reduction in PFC activity, supporting a body of evidence linking traumatic stress with prefrontal hypoactivity.

**Graphical Abstract:** **SPS causes a decrease in excitatory synaptic drive and intrinsic excitability of IL pyramidal neurons.**

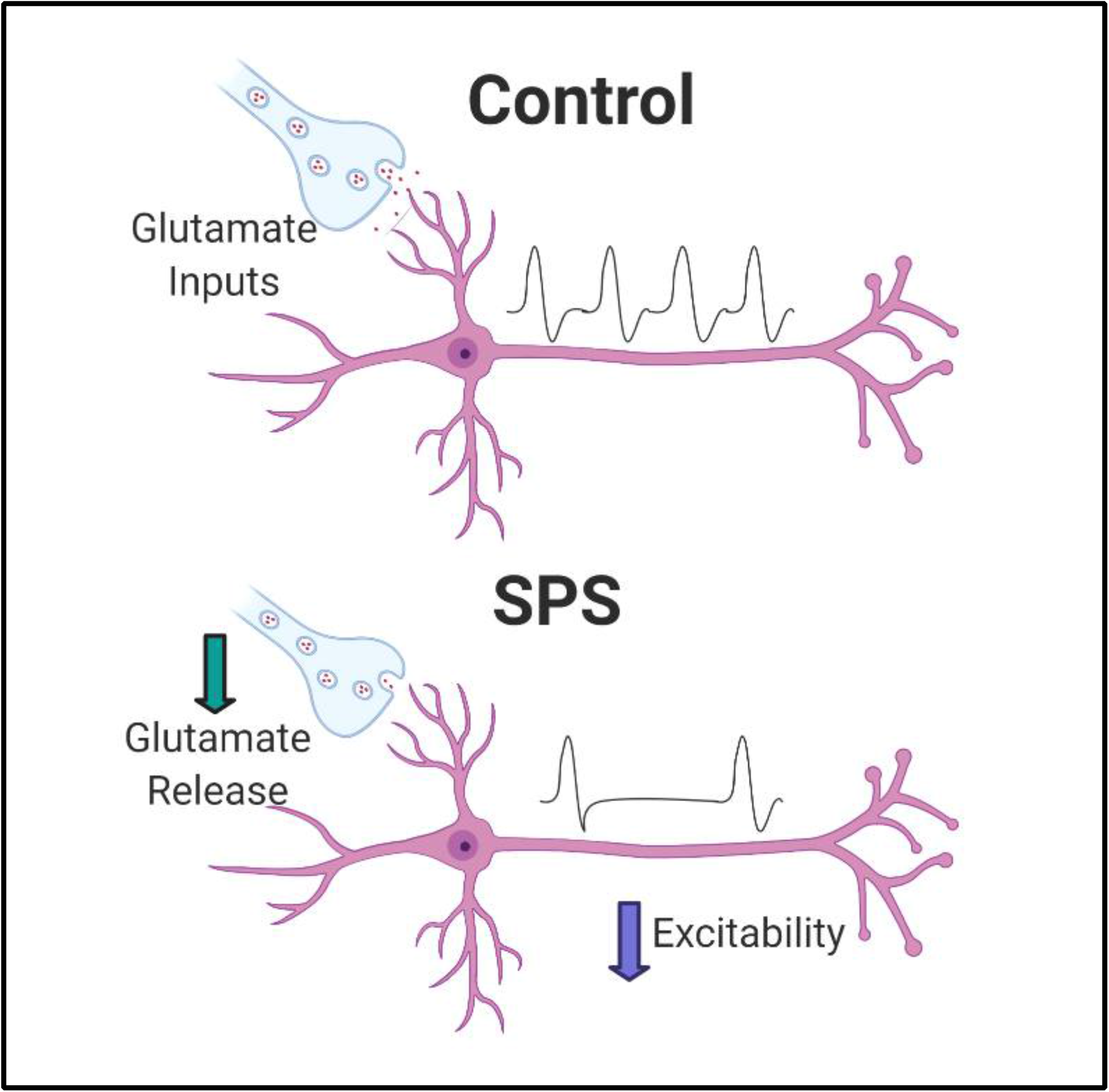

## 1. INTRODUCTION

Posttraumatic stress disorder (PTSD) is among the most prevalent and debilitating neuropsychiatric disorders in the world. In the United States alone, nearly 25 million people will develop PTSD at some point in their lives (Kessler et al., 1995; Maren and Holmes, 2016). To date no universally efficacious treatment exists for PTSD, underscoring the importance of the development of effective therapeutic strategies for this disorder.

Clinical research has linked PTSD with deficits in fear extinction (Milad et al., 2008; Quirk et al., 2006; Ressler et al., 2004), indicative of enhanced responsiveness to emotional stimuli. Symptoms of PTSD can be modeled in rodents using the single prolonged stress (SPS) paradigm (Eagle et al., 2013; Liberzon et al., 1997). Multiple lines of research have shown that SPS disrupts the extinction of fear memories in male rats (Knox et al., 2012; Souza et al., 2017; Yamamoto et al., 2008).

Under normal conditions of fear extinction, excitatory projections from the infralimbic (IL) medial prefrontal cortex (mPFC) to the basolateral amygdala (BLA) activate the intercalated cell clusters, which ultimately reduces output from the central amygdala, thereby promoting extinction of fear (Garcia et al., 1999; Likhtik et al., 2005; Milad and Quirk, 2002; Quirk et al., 2003). However, exposure to an acute traumatic experience may lead to a reduction in activity in the IL mPFC, resulting in a loss of prefrontal inhibition of the amygdala, consistent with exaggerated fear responses (Akirav and Maroun, 2007).

The IL plays an essential role in extinction, extinction recall and reinstatement of conditioned fear, and damage or inactivation of this structure produces extinction memory deficits that resemble those seen in PTSD patients (Norrholm et al., 2011; Sierra-Mercado et al., 2011; Sotres-Bayon and Quirk, 2010). Indeed, abnormally low mPFC activity, together with abnormally high amygdala activity, can be observed in PTSD patients (Liberzon et al., 1999; Milad et al., 2009). Magnetic resonance spectroscopy (MRS) studies indicate that activation of the IL mPFC is reduced following SPS in rats (Piggott et al., 2019), further consistent with a role for reduced IL output in stress pathology. While the role of the IL in promoting fear extinction and emotional regulation is well documented (Kim et al., 2016; Little and Carter, 2013; Milad and Quirk, 2012; Sierra-Mercado et al., 2011), the cellular mechanisms underlying severe stress-related dysfunction remain to be determined.

In this study we tested the impact of SPS on spontaneous excitatory and inhibitory synaptic drive onto principal projection neurons in layer V of the IL as well as their intrinsic membrane excitability. Our results indicate that SPS reduces the intrinsic excitability of IL projection neurons and decreases the efficacy of excitatory signaling onto this population, the latter likely reflecting a reduction in presynaptic glutamate release. We also observed that SPS slowed the decay of GABAergic currents, although this was insufficient to significantly change the overall inhibitory synaptic drive onto IL-PFC neurons. Our results highlight a novel potential mechanism underlying the reduced prefrontal activity observed following SPS, and provides insight into the pathophysiology of abnormal fear memory deficits associated with PTSD.

## 2. METHODS

### 2.1 Rats

Male Sprague Dawley rats were purchased from Envigo and allowed to acclimate for a week at the University of Cincinnati animal housing facility. Rats were maintained under standard conditions (12/12h light/dark cycle, 22 +/- 1°C, food and water *ad libitum*; 2 rats per cage) in accordance with the University of Cincinnati Institutional Animal Care and Use Committee, which specifically approved all stress regimens employed in this study. All animal experiments were carried out in accordance with the National Institutes of Health Guide for the Care and Use of Laboratory Animals (NIH Publications No. 8023, revised 1978). All experiments were performed on adult male rats at 12 weeks of age.

### 2.2 Single Prolonged Stress (SPS) Protocol

Animals were randomly assigned into the control or SPS groups. After the acclimation period, the SPS group was exposed to the SPS paradigm. SPS consisted of three sequential stressors (restraint stress, forced swimming, and ether exposure). First, rats were restrained for 2 h in a plastic animal restrainer, followed immediately by 20 min of forced group swim in water (20–24□°C) in a tub, filled two-thirds from the bottom. Following 15 min of recuperation, rats were exposed to ether (inside a desiccator) until loss of consciousness (less than 5 min) after which rats were placed in their home cages for 7 days without further disturbance (Knox et al., 2012). Control group rats were placed in their home cages for 7 days without any stress.

### 2.3 Electrophysiology

#### 2.3.1 Slice Preparation

Rats were sacrificed 7 days post SPS at approximately postnatal day 91. The 7 day incubation time was selected as behavioral abnormalities are consistently observed following this incubation period (Iwamoto et al., 2007; Keller et al., 2015; Knox et al., 2016, 2012). Animals were deeply anesthetized with sodium pentobarbital (390 mg/kg, Fatal-Plus) and decapitated. A warm slicing protocol was used to prepare healthy adult rat brain slices as previously described (Ting et al., 2014). Brains were quickly isolated and dura matter carefully removed before removing the cerebellum. The brain was then immediately glued to a cutting stage and immersed in NMDG solution (92 mM NMDG, 2.5 mM KCl, 1.2 mM NaH_2_PO_4_, 30 mM NaHCO_3_, 20 mM HEPES, 25 mM glucose, 5 mM sodium ascorbate, 2 mM thiourea, 3 mM sodium pyruvate, 10 mM MgSO_4_, and 0.5 mM CaCl_2_) at a temperature of 34-36°C and continuously bubbled with 95% oxygen and 5% carbon-dioxide. Coronal slices containing the mPFC were sectioned at 300 μm thickness using a vibrating microtome (Vibratome 7000smz-2; Campden Instruments, Lafayette, IN) with ceramic blades (Campden Instruments) at an advance speed of 0.03 mm/s. Vertical vibration of the blade was manually tuned in accordance with the user manual, and was set to 0.1 – 0.3 μm. Bath temperature was kept within the desired range of 34-36°C, by adding warm or cold water into the external chamber of the Vibratome, and was monitored throughout the cutting procedure with a conventional mercury/glass thermometer. The slices were allowed to recover for 1 hour in oxygenated NMDG solution at 34-36°C. At the end of recovery, slices were transferred to a chamber containing oxygenated artificial CSF solution (125 mM NaCl, 2.5 mM KCl, 25 mM NaHCO_3_, 1 mM NaH_2_PO_4_, 25 mM glucose, 1 mM MgCl_2_, 2 mM CaCl_2_) for at least 30 minutes at room temperature after which the slices were ready for in vitro patch clamp recordings.

#### 2.3.2 Electrophysiological recording

Brain slices were transferred to a submersion-type recording chamber (RC-22; Warner Instruments, Hamden, CT) and mounted onto the stage of an upright microscope (BX51WI, Olympus, Center Valley, PA). Slices were then perfused at a flow rate of 2–4 ml/min with oxygenated aCSF at 34-36°C. Patch electrodes were constructed from thin-walled single-filamented borosilicate glass (1.5 mm outer diameter; World Precision Instruments) using a microelectrode puller (P-97; Sutter Instruments, Novato, CA). Pipette resistances ranged from 4 to 6 MΩ, and seal resistances were > 1 GΩ.

Whole-cell patch clamp recordings were obtained from layer V pyramidal neurons in the mPFC using a MultiClamp 700B amplifier (Molecular Devices, Sunnyvale, CA). Pyramidal neurons were easily identifiable in the slice based on soma morphology and the presence of a prominent apical dendrite. For all electrophysiological recordings, membrane voltages were adjusted for liquid junction potentials (approximately –14 mV) calculated using JPCalc software (P. Barry, University of New South Wales, Sydney, Australia; modified for Molecular Devices). Signals were filtered at 4–6 kHz through a –3 dB, four-pole low-pass Bessel filter and digitally sampled at 20 kHz using a commercially available data acquisition system (Digidata 1550A with pClamp 10.5 software; Molecular Devices). Data were recorded using pClamp and stored on a computer for offline analysis. Current clamp recordings were analyzed using Clampfit (Molecular Devices). For studies examining synaptic transmission, the amplitude and frequency of miniature excitatory postsynaptic currents (mEPSCs) and miniature inhibitory postsynaptic currents (mIPSCs) were measured using MiniAnalysis 6.0.7 (Synaptosoft; Decatur, GA, USA), and the threshold for mEPSC and mIPSC detection was set at twice the root mean square (RMS) of the background noise.

#### 2.3.3 Intrinsic Excitability Measurements

For intrinsic excitability measurements, patch electrodes were filled with a solution containing the following: 130 mM K-gluconate, 10 mM KCl, 10 mM HEPES, 10 mM sodium phosphocreatine, 4 mM MgATP, and 0.3 mM Na_2_-GTP, pH 7.2, 295-300 mOsm. In the current clamp mode, once a stable membrane potential was observed, intrinsic excitability measurements were performed at the resting membrane potential (RMP) approximately 1 min after whole-cell configuration was established. Cell capacitance was measured using the membrane test function in pClamp 10.5 (Molecular Devices, Sunnyvale, CA, USA). All measurements of intrinsic membrane excitability were taken from RMP. Rheobase was measured by applying depolarizing current steps (10 pA steps, 100 msec duration) until the generation of a single action potential (AP). Input (membrane) resistance (R_input_) was measured by applying a hyperpolarizing current step (−10 pA) via the patch electrode. AP threshold was defined as the *V*_m_ measured 0.5 ms before the peak in the second derivative of the waveform. The action potential threshold and amplitude were analyzed for the first spike at the rheobase current injection. AP half-width (AP_50_) was determined by measuring the elapsed time from the peak of the AP to 50% maximum amplitude during the repolarization phase. Firing rate was measured in response to 20 pA depolarizing current steps in the current clamp configuration. The number of action potentials generated over a period of 1 second was recorded across the stimulus intensity range of 0-280 pA. All intrinsic excitability measurements were conducted in oxygenated aCSF at 34-36°C. Cells with RMP lower than −55mV were included in the final analysis. Recordings were obtained from 14-17 cells from 3 rats in each group.

#### 2.3.4 Synaptic Drive Measurements

To measure synaptic drive, miniature postsynaptic currents (mPSCs) were recorded in the presence of TTX (0.5□μM; Hellobio; Princeton, NJ, USA). Patch electrodes were filled with a solution containing the following (in mM): 130 Cs-gluconate, 10 CsCl, 10 HEPES, 11 EGTA, 1 CaCl_2_, and 2 MgATP, pH 7.2 (295–305□mOsm). In order to isolate mEPSCs, cells were voltage clamped at −70□mV. To record mIPSCs, cells were held at 0 mV. Peak mPSC amplitude was measured from baseline. Decay kinetics were estimated using a single exponential function: [y(t)=a*exp(-t/τ)] using the average mPSC in a given neuron. Synaptic drive was measured in each sampled neuron by multiplying the area under the average mPSC by the mPSC frequency. mEPSC and mIPSC recordings were obtained from the same 14-18 cells from three rats in each group. For mIPSC recordings, one additional animal was included to obtain a total of 23-27 cells in each group.

#### 2.3.5 Statistical Analysis

All data sets were tested for normality using the Kolmogorov-Smirnoff test. Data were analyzed by unpaired t-test when groups were normally distributed. The Mann-Whitney non-parametric test was performed for groups not following a normal distribution. AP firing rate was analyzed by two-way repeated measures ANOVA with SPS and stimulus intensity as factors. In the cases where significant differences and interactions were found, multiple comparisons with false discovery rate correction (FDR) was performed for post hoc analysis. Data were analyzed using Prism 8 (GraphPad Software, La Jolla California). Outliers for normally distributed dataset were calculated using Prism Grubbs’ test and excluded from the analysis.

## 3. RESULTS

### 3.1 SPS Reduces the Intrinsic Excitability of Layer V Pyramidal Neurons in the IL mPFC

One week following SPS, patch clamp recordings were obtained from layer V pyramidal neurons under the current clamp configuration to probe potential changes in the intrinsic membrane properties of this population (Figure 1A). Rheobase was increased in animals exposed to SPS (t= 5.6, df=31, p<0.0001, n=16 cells from control and SPS respectively; Figure 1B). SPS also decreased input (membrane) resistance (U=29, p=0.001, n=14 cells from control and SPS respectively; Figure 1C), which may contribute to the increase in rheobase observed following SPS. There was a significant decrease in RMP following SPS (U=45, p=0.001, n=16 cells from control and SPS respectively; Figure 1F). SPS also significantly decreased AP_50_ (t= 3.8, df=30, p=0.0006, n=16 cells from control and SPS respectively; Figure 1G). Mean capacitance of the cells were 90.2 +/- 3.4 pF and 94.8+/- 5.6 pF for control and SPS groups respectively. No significant change in AP threshold (t= 0.55, df=30, p=0.6, n=16 cells from control and SPS respectively; Figure 1D) or AP amplitude (t=0.96, df=29, p=0.34, n=16 and 15 cells from control and SPS respectively; Figure 1E) were observed following SPS.

**Fig. 1.**
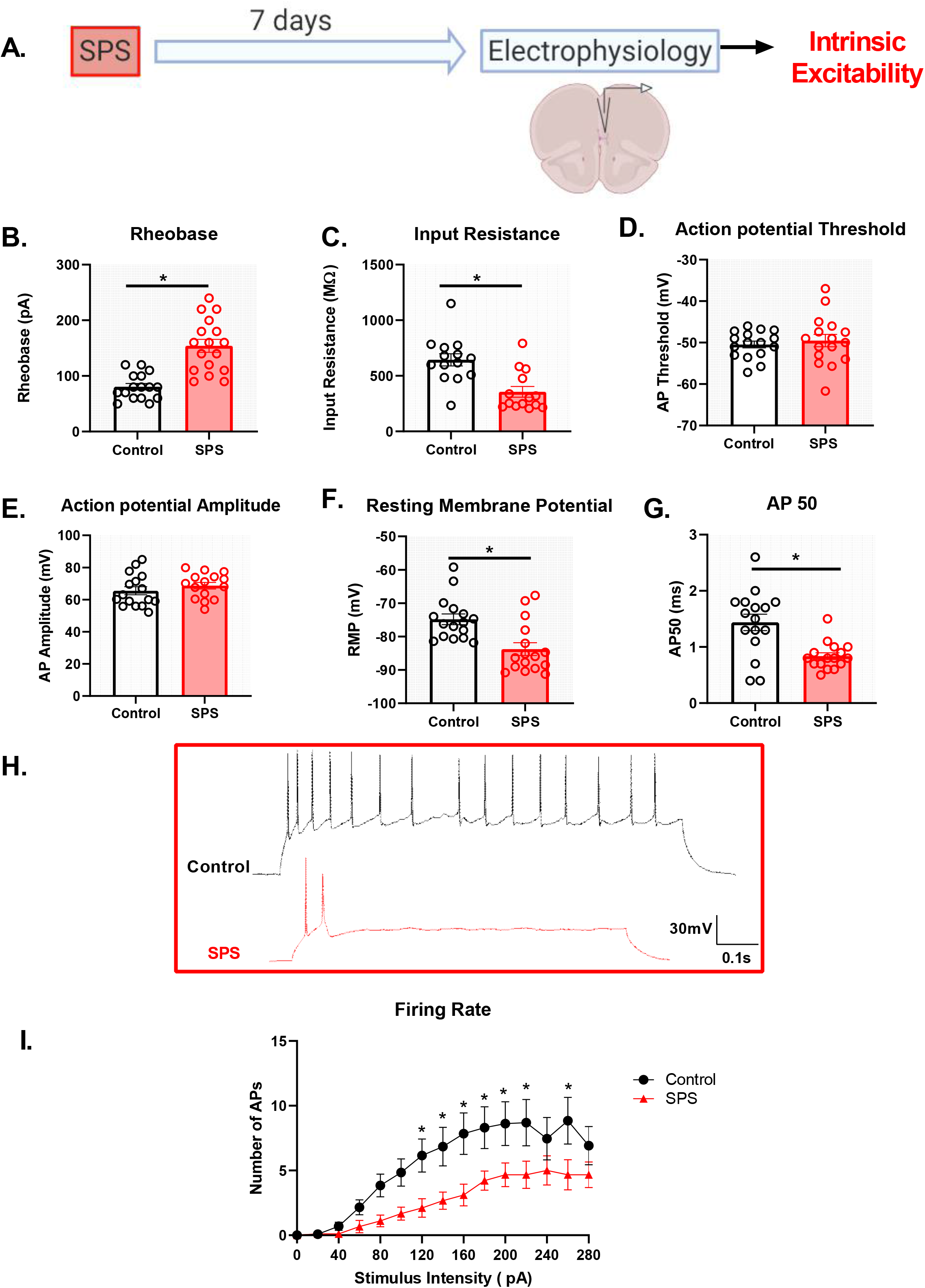
SPS decreases the intrinsic excitability of IL pyramidal neurons. Schematic of the experimental timeline (A). Electrophysiological recordings in current clamp mode from the IL mPFC were conducted in male rats 7 days post SPS. SPS increases rheobase (B) (t(31)=5.6, p<0.0001), decreases input resistance (C) (Mann-Whitney U(26)=29, p<0.01), hyperpolarizes RMP (F) (Mann-Whitney U(34)=45, p<0.01), decreases AP_50_ (G) (t(30)=3.8, p<0.001) and also decreases firing rate of IL pyramidal neurons (I) with a main effect of SPS [F(1,20)=4.7; p<0.05], main effect of stimulus intensity [F(14,280)=20.3; p<0.01] and a significant SPS X stimulus intensity interaction [F(14,280)=2.03; p<0.01]. The SPS group had a significantly lower action potential firing rate compared to controls at a stimulus intensity range of 120-220 pA and at 260 pA (p<0.05). Representative traces of action potentials in control (black) vs SPS (red) groups following 160 pA current injection are shown in (H). Scale bars: 30mV, 0.1s. SPS had no effect on action potential threshold (D) (t(30)=0.55, p>0.5) or amplitude (E) (t(29)=0.96, p>0.5). Data presented as Mean +/- SEM.

We next analyzed the repetitive firing rate of the pyramidal neurons following SPS (Figure 1H and I). There was a significant SPS X stimulus intensity interaction [F(14,280)=2.03; p=0.002, n=13 and 9 cells from control and SPS respectively; Figure 1I]. Multiple comparisons with FDR correction indicate that the SPS group had significantly lower action potential firing compared to the control group at a stimulus intensity range of 120-220 pA and at 260 pA (p<0.05). Collectively, these results suggest that SPS reduces the intrinsic membrane excitability of layer V IL pyramidal neurons.

### 3.2 SPS Reduces Excitatory Synaptic Drive onto Layer V Pyramidal Neurons in the IL mPFC

Previous studies indicate reduced overall glutamate levels in the mPFC following SPS (Piggott et al., 2019), but the underlying mechanisms by which SPS alters glutamatergic transmission in the region remain unclear. Seven days after SPS, mEPSCs were recorded in pyramidal neurons under voltage clamp conditions at holding potential of −70mV (Figure 2A). Our results show that SPS significantly reduces the frequency of mEPSCs (t= 3.9, df=32, p=0.0004, n=18 and 16 cells from control and SPS respectively; Figure 2B, C) while having no effect on mEPSC amplitude (t= 0.9, df=31, p=0.33, n=17 and 16 cells from control and SPS respectively; Figure 2D) or mEPSC decay (t=0.1, df=29, p=0.91, n=17 and 14 cells from control and SPS respectively; Figure 2E). Finally, we show that SPS decreases overall excitatory synaptic drive onto pyramidal neurons in the IL mPFC (Mann-Whitney U=62, p=0.03, n=16 cells from control and SPS respectively; Figure 2F). Collectively these data suggest that SPS reduces spontaneous excitatory synaptic drive onto IL layer V pyramidal neurons, which is likely driven by a reduction in the presynaptic release of glutamate.

**Fig. 2.**
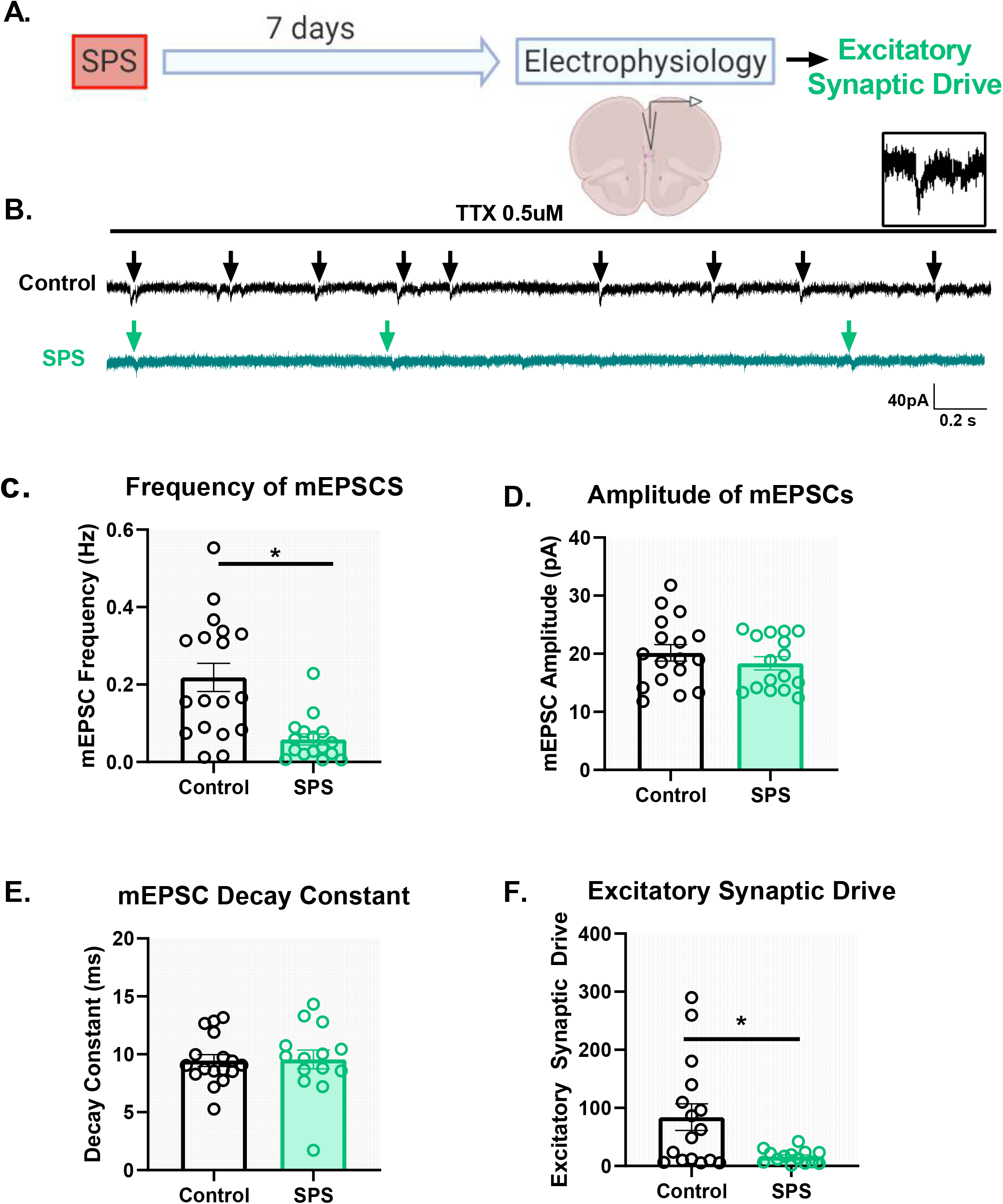
SPS decreases spontaneous glutamatergic drive onto IL pyramidal neurons. Schematic of the experimental timeline (A). Electrophysiological recordings in voltage clamp mode from the IL mPFC were conducted in male rats 7 days post SPS. Representative voltage clamp mEPSC traces of control (black) and SPS (green) groups are shown in (B). Arrows indicate mEPSC events. Scale bars: 40pA, 0.2s. Magnified image of a single mEPSC event is shown on top right (B). SPS decreases frequency of mEPSCs (C) (t(32)=3.9, p<0.01). SPS has no effect on mEPSC amplitude (D) (t(32)=0.9, p=0.3) or mEPSC decay rate (E) (t(29)=0.1, p=0.9). SPS significantly decreases excitatory synaptic drive (F) (Mann-Whitney U=62, p<0.05). Data presented as Mean +/- SEM.

### 3.3 SPS Prolongs the Decay of GABA Currents with No Effect on Total Inhibitory Synaptic Drive onto Layer V IL mPFC Pyramidal Neurons

Similar to the excitatory synaptic drive experiments, mIPSCs were measured 7 days following SPS under the voltage clamp configuration at a holding potential of 0 mV (Figure 3A). Analysis of mIPSC frequency (t= 1.62, df=49, p=0.11, n=24 and 27 cells from control and SPS respectively; Figure 3B, C) and amplitude (t=2, df=48, p=0.05, n=23 and 27 cells from control and SPS respectively; Figure 3D) did not reveal any significant effects. However, analysis of the mIPSC decay constant showed that SPS significantly prolongs the decay of GABA currents in the IL mPFC (t=3.5, df=48, p=0.001, n=24 and 26 cells from control and SPS respectively; Figure 3E).

**Fig. 3.**
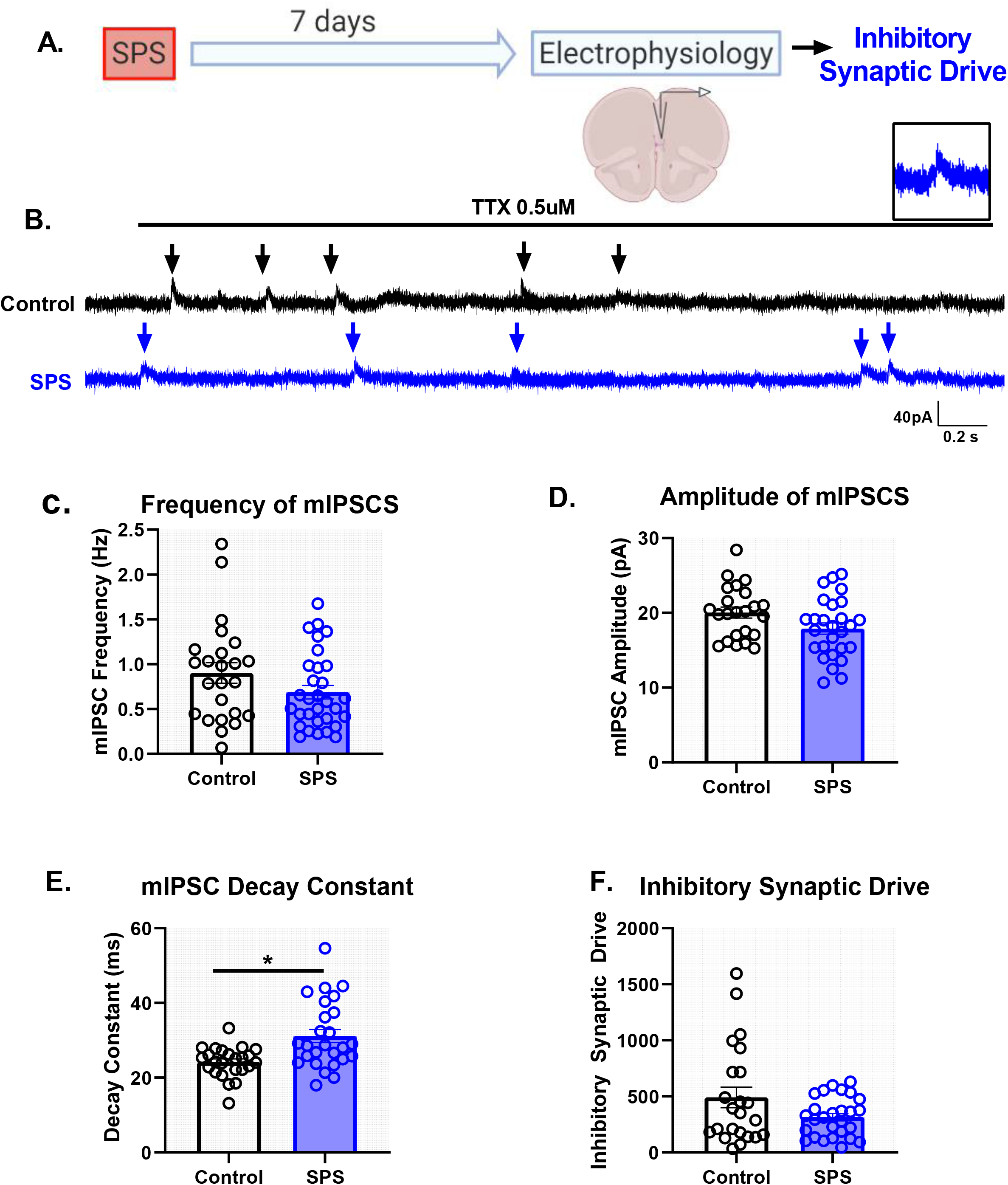
SPS prolongs the decay of GABA currents but has no effect on overall spontaneous inhibitory synaptic drive in the IL. Schematic of the experimental timeline (A). Electrophysiological recordings in the voltage clamp mode were obtained from the IL mPFC in male rats 7 days post SPS. Representative voltage clamp mIPSC traces of control (black) and SPS (blue) groups are shown in (B). Arrows indicate mIPSC events. Scale bars: 40pA, 0.2s. Magnified image of a single mIPSC event is shown on top right (B). SPS has no effect on mIPSC frequency (C) (t(49)=1.6, p=0.1) or mIPSC amplitude (D) (t(48)=2.0, p=0.05). SPS increases the mIPSC decay (E) (t(48)=3.5, p<0.01) but has no effect on inhibitory synaptic drive (F) (Mann-Whitney U=263, p=0.3). Data presented as Mean +/- SEM.

Nonetheless, analysis of total inhibitory synaptic drive did not reveal any significant difference between the SPS and Control groups (Mann-Whitney U=263, p=0.3, n=23 and 27 cells from control and SPS respectively; Figure 3F). Collectively, these data suggest that SPS might not affect the presynaptic release of GABA in the IL, but may allow for GABA to be present in the synaptic cleft longer as demonstrated by the reduction in the decay of GABA currents. However, that effect does not result in significant changes in the strength of spontaneous synaptic inhibition within the IL following SPS.

## 4. DISCUSSION

Our results indicate that SPS causes physiological changes in IL mPFC glutamatergic pyramidal neurons and their associated synaptic inputs. Here we demonstrate that SPS reduces the intrinsic membrane excitability of IL pyramidal neurons, as indicated by an increased rheobase, decreased input resistance, hyperpolarized RMP, and a reduction in repetitive firing rate. In addition, SPS causes alterations in spontaneous synaptic drive onto the major output pyramidal neurons in layer V of the IL. SPS reduces the excitatory glutamatergic synaptic tone driven mainly by decreases in presynaptic inputs. Our results further indicate that SPS prolongs the decay of GABA currents in the IL but does not change total inhibitory synaptic drive. Collectively, these data suggest possible mechanisms via which SPS may cause a reduction in IL mPFC activity and contribute towards a better understanding of the pathophysiology associated with PTSD symptoms.

SPS causes deficits in fear learning and extinction of fear responses (Iwamoto et al., 2007; Keller et al., 2015; Knox et al., 2016, 2012). The IL mPFC in rodents and the ventromedial cortex (Brodmann area 25) in humans plays a role in driving extinction of fear responses (Milad and Quirk, 2012; Quirk et al., 2000; Quirk and Mueller, 2008). Prefrontal hypoactivity and subsequent amygdala hyperactivity leads to a disruption of the top-down executive control of fear responses, and is thought to underlie the abnormal extinction of fear responses in human PTSD patients and also in rodent models of traumatic stress (Milad et al., 2009; Piggott et al., 2019; Quirk et al., 2003). Prior studies indicate that SPS results in reduced Fos activation in IL mPFC suggesting reduced prefrontal drive following SPS, which might underlie the abnormal fear extinction observed in SPS treated animals (Piggott et al., 2019). Our current clamp results indicate that following SPS, the ability to generate an action potential is impaired in IL layer V pyramidal cells (as evidenced by the increased rheobase, reduced input resistance and hyperpolarized RMP). Furthermore, once firing is initiated, SPS animals show a reduction in the number of action potentials across different stimulus intensities compared to controls. Our findings suggest that SPS reduces the intrinsic membrane excitability of glutamatergic pyramidal neurons in the IL mPFC.

Our voltage clamp experiments show that SPS reduces spontaneous excitatory synaptic drive onto IL pyramidal neurons. Specifically, our findings indicate that SPS reduces the frequency of mEPSCs without causing any change in mEPSC amplitude. These results suggest that SPS causes a reduction in presynaptic glutamate release onto pyramidal neurons without affecting postsynaptic NMDA/AMPA receptor function (Liley, 1956). The changes in presynaptic input might be due to changes in the probability of glutamate release or the number of glutamatergic synaptic contacts onto the pyramidal neurons, and further studies are needed to answer that question. Nevertheless, our results are consistent with previous reports showing reduced glutamate levels in the mPFC following SPS (Piggott et al., 2019) and further suggest that the stress-evoked reduction in glutamatergic signaling within the IL cortex may be presynaptically mediated.

It is not known which long-range excitatory inputs, or pyramidal neuron outputs, are specifically affected by SPS. Afferent input to the IL includes glutamatergic input from regions implicated in emotional memory, including the BLA, thalamus and ventral hippocampus (vHPC) (Hoover and Vertes, 2007). Importantly, BLA-IL projections are selectively activated during extinction of conditioned fear, and stimulation of the BLA-IL connection facilitates extinction of conditioned fear (Senn et al., 2014). Additionally, SPS selectively affects the IL-BLA projection and does not affect the IL-dorsolateral striatum projection (Piggott et al., 2019). IL-BLA projections are important for extinction of fear responses (Bloodgood et al., 2018). The IL also projects to the periaqueductal grey (PAG), a circuit that is also known to be involved in fear responses (Cheriyan et al., 2016). Taking all the above findings, it is possible that SPS might target BLA-IL-PAG or reciprocal BLA-IL-BLA connections. Future studies should aim to explore this potential circuit specificity of the physiological effects following SPS.

Processes underlying SPS-induced decreases in IL intrinsic excitability or excitatory synaptic drive remain to be determined. The increase in rheobase, decrease in input resistance and a more hyperpolarized RMP indicates that it is more difficult to depolarize the neuron to spike threshold following SPS. A more hyperpolarized RMP and reduced membrane resistance after SPS could be due to an increase in the number, or conductance, of “leak” K^+^ channels, resulting in a greater K+ efflux from the cells and reduced excitability (Honoré, 2007). Rat mPFC pyramidal neurons also express KCNQ2 channels (K_v_7 voltage gated K+ channel), the excessive opening of which might reduce neuronal firing in the PFC following stress (Arnsten et al., 2019). Some forms of KCNQ channels are constitutively active and may contribute towards the leak conductance (Goldstein et al., 2001; Schroeder et al., 2000).

Several lines of evidence also implicate G-protein gated inwardly rectifying K^+^ (GIRK) channel dysfunction in stress-related alterations in the excitability of prefrontal neurons and psychiatric disorders (Clarke et al., 2011; Victoria et al., 2016). Activation of GIRK channels results in K^+^ efflux, which hyperpolarizes the neuronal RMP and dampens neuronal excitability (Dascal, 1997; Hibino et al., 2010). Acute stress can increase NMDA and AMPAR-mediated synaptic currents in PFC (Yuen et al., 2009), which can increase the expression of GIRK channels (Hee et al., 2009). Thus, the decreased excitability observed following SPS could be due to an increase in expression of GIRK channels, which might be a compensation mechanism for the increase in excitatory neurotransmission that is typically observed immediately following acute stress. It is also possible that SPS may increase constitutive GIRK channel activity (Chen and Johnston, 2005; Takigawa and Alzheimer, 2002), leading to membrane hyperpolarization and a reduction in excitability. Importantly, mouse models exhibiting enhanced GIRK-dependent signaling show deficits in fear learning (Cooper et al., 2012) suggesting that an increase in GIRK channel signaling and reduction in neuronal excitability might also underlie aberrant fear responses observed following SPS. Further studies will be needed to test this hypothesis.

Literature evidence regarding changes in GABAergic signaling following SPS is inconsistent. Some studies using MRS show no change in GABA levels following SPS (Knox et al., 2010; Piggott et al., 2019) in rodents or in humans with PTSD (Rosso et al., 2014; Schür et al., 2016). Our results demonstrate a slower decay of GABA currents following SPS, with no evidence for changes in presynaptic GABA release or postsynaptic GABA receptor expression. Slower GABA decay could be due to factors such as reduced neurotransmitter uptake and delayed clearance (Overstreet and Westbrook, 2003), slower deactivation time of GABA-A receptors (Schofield and Huguenard, 2007) and changes in GABA-A receptor subunit composition which may alter postsynaptic GABA channel closing kinetics (Haas and Macdonald, 1999). Benzodiazepines can also prolong the decay of GABA currents by changing conformation of the GABA-A receptor such that it slows GABA current deactivation (Bianchi et al., 2007; Xiong et al., 2018). Thus, the prolonged GABA decay we observed following SPS might also be due to alteration in benzodiazepine binding leading to slower receptor deactivation and longer decay.

In this study we recorded from layer V pyramidal neurons identified by a thick apical dendrite. Prior evidence suggests that pyramidal neurons with thick tufted apical dendrites belong to the Type A class, which receive strong GABAergic innervation from fast spiking parvalbumin (PV) interneurons (INs) (Lee et al., 2014). Therefore, it is possible that SPS-mediated plasticity was only occurring in specific microcircuits such as the PV-to-pyramidal neuron connection. Indeed, prior studies indicate that neurons projecting from IL to BLA receive strong innervation via PV INs (Marek et al., 2018).

Therefore, it will be important to explore how SPS alters inhibitory synaptic connections between PV INs and the IL-PFC pyramidal neurons projecting to the BLA or PAG.

Taken together, our findings suggest that increasing the intrinsic excitability and glutamatergic synaptic input onto IL pyramidal neurons might be effective in preventing some of the behavioral changes observed with SPS. Indeed, various studies indicate that enhanced top-down control of subcortical regions leads to more efficient control of emotion regulation. fMRI studies in humans have shown that greater prefrontal drive may be a resilience factor in PTSD (Chen et al., 2018). Recent studies indicate increased neuronal activation of mPFC in resilient mice following chronic predator or social defeat stress (Adamec et al., 2012). Consistent with this hypothesis, direct optogenetic stimulation of the ventral portion of the mPFC has been shown to promote resilience to social defeat stress (Covington et al., 2010).

## 5. CONCLUSION

Overall, our findings indicate that reduced prefrontal drive following SPS may underlie the abnormal fear responses observed with the stress paradigm. Our results highlight novel multifaceted mechanisms by which SPS can cause a reduction in PFC activity, supporting growing evidence that severe stress leads to prefrontal hypoactivity, a characteristic of diseases such as PTSD.

## Authors contribution

**NN,MB,JH:** Conceptualization, Methodology, Software. **NN:** Data collection, analysis, Writing-Original draft preparation. **MB, JH**: Supervision. **NN,MB, JH:** Writing-Reviewing and Editing

## Acknowledgements

The authors would like to thank Ana Franco-Villanueva and Kristen Oshima for guidance in electrophysiology and help with recordings. Images for this paper were created with BioRender.com

## References

Adamec, R., Toth, M., Haller, J., Halasz, J., Blundell, J., 2012. A comparison of activation patterns of cells in selected prefrontal cortical and amygdala areas of rats which are more or less anxious in response to predator exposure or submersion stress. Physiol. Behav. 105, 628–638. https://doi.org/10.1016/J.PHYSBEH.2011.09.016

Akirav, I., Maroun, M., 2007. The role of the medial prefrontal cortex-amygdala circuit in stress effects on the extinction of fear. Neural Plast. 2007, 30873. https://doi.org/10.1155/2007/30873

Arnsten, A.F.T., Jin, L.E., Gamo, N.J., Ramos, B., Paspalas, C.D., Morozov, Y.M., Kata, A., Bamford, N.S., Yeckel, M.F., Kaczmarek, L.K., El-Hassar, L., 2019. Role of KCNQ potassium channels in stress-induced deficit of working memory. Neurobiol. Stress 11. https://doi.org/10.1016/j.ynstr.2019.100187

Bianchi, M.T., Botzolakis, E.J., Haas, K.F., Fisher, J.L., Macdonald, R.L., 2007. Microscopic kinetic determinants of macroscopic currents: Insights from coupling and uncoupling of GABAA receptor desensitization and deactivation. J. Physiol. 584, 769–787. https://doi.org/10.1113/jphysiol.2007.142364

Bloodgood, D.W., Sugam, J.A., Holmes, A., Kash, T.L., 2018. Fear extinction requires infralimbic cortex projections to the basolateral amygdala. Transl. Psychiatry 8. https://doi.org/10.1038/s41398-018-0106-x

Chen, F., Ke, J., Qi, R., Xu, Q., Zhong, Y., Liu, T., Li, J., Zhang, L., Lu, G., 2018. Increased Inhibition of the Amygdala by the mPFC may Reflect a Resilience Factor in Post-traumatic Stress Disorder: A Resting-State fMRI Granger Causality Analysis. Front. Psychiatry 9, 516. https://doi.org/10.3389/fpsyt.2018.00516

Cheriyan, J., Kaushik, M.K., Ferreira, A.N., Sheets, P.L., 2016. Specific Targeting of the Basolateral Amygdala to Projectionally Defined Pyramidal Neurons in Prelimbic and Infralimbic Cortex. eNeuro 3. https://doi.org/10.1523/ENEURO.0002-16.2016

Clarke, T.K., Laucht, M., Ridinger, M., Wodarz, N., Rietschel, M., Maier, W., Lathrop, M., Lourdusamy, A., Zimmermann, U.S., Desrivieres, S., Schumann, G., 2011. KCNJ6 is associated with adult alcohol dependence and involved in gene × early life stress interactions in adolescent alcohol drinking. Neuropsychopharmacology 36, 1142–1148. https://doi.org/10.1038/npp.2010.247

Cooper, A., Grigoryan, G., Guy-David, L., Tsoory, M.M., Chen, A., Reuveny, E., 2012. Trisomy of the G protein-coupled K + channel gene, Kcnj6, affects reward mechanisms, cognitive functions, and synaptic plasticity in mice. Proc. Natl. Acad. Sci. U. S. A. 109, 2642–2647. https://doi.org/10.1073/pnas.1109099109

Covington, H.E., Lobo, M.K., Maze, I., Vialou, V., Hyman, J.M., Zaman, S., LaPlant, Q., Mouzon, E., Ghose, S., Tamminga, C.A., Neve, R.L., Deisseroth, K., Nestler, E.J., 2010. Antidepressant Effect of Optogenetic Stimulation of the Medial Prefrontal Cortex. J. Neurosci. 30, 16082–16090. https://doi.org/10.1523/JNEUROSCI.1731-10.2010

Dascal, N., 1997. Signalling via the G protein-activated K+ channels. Cell. Signal. 9, 551–573. https://doi.org/10.1016/S0898-6568(97)00095-8

Eagle, A.L., Knox, D., Roberts, M.M., Mulo, K., Liberzon, I., Galloway, M.P., Perrine, S.A., 2013. Single prolonged stress enhances hippocampal glucocorticoid receptor and phosphorylated protein kinase B levels. Neurosci. Res. 75, 130–137. https://doi.org/10.1016/j.neures.2012.11.001

Garcia, R., Vouimba, R.-M., Baudry, M., Thompson, R.F., 1999. The amygdala modulates prefrontal cortex activity relative to conditioned fear. Nature 402, 294–296w. https://doi.org/10.1038/46286

Goldstein, S.A.N., Bockenhauer, D., O’Kelly, I., Zilberberg, N., 2001. Potassium leak channels and the KCNK family of two-p-domain subunits. Nat. Rev. Neurosci. 2, 175–184. https://doi.org/10.1038/35058574

Haas, K.F., Macdonald, R.L., 1999. GABA(A) receptor subunit γ2 and δ subtypes confer unique kinetic properties on recombinant GABA(A) receptor currents in mouse fibroblasts. J. Physiol. 514, 27–45. https://doi.org/10.1111/j.1469-7793.1999.027af.x

Hee, J.C., Qian, X., Ehlers, M., Yuh, N.J., Jan, L.Y., 2009. Neuronal activity regulates phosphorylation-dependent surface delivery of G protein-activated inwardly rectifying potassium channels. Proc. Natl. Acad. Sci. U. S. A. 106, 629–634. https://doi.org/10.1073/pnas.0811615106

Hibino, H., Inanobe, A., Furutani, K., Murakami, S., Findlay, I., Kurachi, Y., 2010. Inwardly rectifying potassium channels: Their structure, function, and physiological roles. Physiol. Rev. https://doi.org/10.1152/physrev.00021.2009

Hoover, W.B., Vertes, R.P., 2007. Anatomical analysis of afferent projections to the medial prefrontal cortex in the rat. Brain Struct. Funct. 212, 149–179. https://doi.org/10.1007/s00429-007-0150-4

Iwamoto, Y., Morinobu, S., Takahashi, T., Yamawaki, S., 2007. Single prolonged stress increases contextual freezing and the expression of glycine transporter 1 and vesicle-associated membrane protein 2 mRNA in the hippocampus of rats. Prog. Neuro-Psychopharmacology Biol. Psychiatry 31, 642–651. https://doi.org/10.1016/j.pnpbp.2006.12.010

Keller, S.M., Schreiber, W.B., Stanfield, B.R., Knox, D., 2015. Inhibiting corticosterone synthesis during fear memory formation exacerbates cued fear extinction memory deficits within the single prolonged stress model. Behav. Brain Res. 287, 182–186. https://doi.org/10.1016/j.bbr.2015.03.043

Kessler, R.C., Sonnega, A., Bromet, E., Hughes, M., Nelson, C.B., 1995. Posttraumatic Stress Disorder in the National Comorbidity Survey. Arch. Gen. Psychiatry 52, 1048–1060. https://doi.org/10.1001/archpsyc.1995.03950240066012

Kim, H.S., Cho, H.Y., Augustine, G.J., Han, J.H., 2016. Selective Control of Fear Expression by Optogenetic Manipulation of Infralimbic Cortex after Extinction. Neuropsychopharmacology 41, 1261–1273. https://doi.org/10.1038/npp.2015.276

Knox, D., George, S.A., Fitzpatrick, C.J., Rabinak, C.A., Maren, S., Liberzon, I., 2012. Single prolonged stress disrupts retention of extinguished fear in rats. Learn. Mem. 19, 43–9. https://doi.org/10.1101/lm.024356.111

Knox, D., Perrine, S.A., George, S.A., Galloway, M.P., Liberzon, I., 2010. Single prolonged stress decreases glutamate, glutamine, and creatine concentrations in the rat medial prefrontal cortex. Neurosci. Lett. 480, 16–20. https://doi.org/10.1016/j.neulet.2010.05.052

Knox, D., Stanfield, B.R., Staib, J.M., David, N.P., Keller, S.M., DePietro, T., 2016. Neural circuits via which Single prolonged stress exposure leads to fear extinction retention deficits. Learn. Mem. 23, 689–698. https://doi.org/10.1101/lm.043141.116

Lee, A.T., Gee, S.M., Vogt, D., Patel, T., Rubenstein, J.L., Sohal, V.S., 2014. Pyramidal Neurons in Prefrontal Cortex Receive Subtype-Specific Forms of Excitation and Inhibition. https://doi.org/10.1016/j.neuron.2013.10.031

Liberzon, I., Krstov, M., Young, E.A., 1997. Stress-restress: Effects on ACTH and fast feedback. Psychoneuroendocrinology 22, 443–453. https://doi.org/10.1016/S0306-4530(97)00044-9

Liberzon, I., Taylor, S.F., Amdur, R., Jung, T.D., Chamberlain, K.R., Minoshima, S., Koeppe, R.A., Fig, L.M., 1999. Brain activation in PTSD in response to trauma-related stimuli. Biol. Psychiatry 45, 817–826. https://doi.org/10.1016/S0006-3223(98)00246-7

Likhtik, E., Pelletier, J.G., Paz, R., Paré, D., 2005. Prefrontal Control of the Amygdala. J. Neurosci. 25, 7429–7437. https://doi.org/10.1523/JNEUROSCI.2314-05.2005

Liley, A.W., 1956. The quantal components of the mammalian end-plate potential. J. Physiol. 133, 571–587. https://doi.org/10.1113/jphysiol.1956.sp005610

Little, J.P., Carter, A.G., 2013. Synaptic mechanisms underlying strong reciprocal connectivity between the medial prefrontal cortex and basolateral amygdala. J. Neurosci. 33, 15333–15342. https://doi.org/10.1523/JNEUROSCI.2385-13.2013

Marek, R., Jin, J., Goode, T.D., Giustino, T.F., Wang, Q., Acca, G.M., Holehonnur, R., Ploski, J.E., Fitzgerald, P.J., Lynagh, T., Lynch, J.W., Maren, S., Sah, P., 2018. Hippocampus-driven feed-forward inhibition of the prefrontal cortex mediates relapse of extinguished fear. Nat. Neurosci. 21, 384–392. https://doi.org/10.1038/s41593-018-0073-9

Maren, S., Holmes, A., 2016. Stress and fear extinction. Neuropsychopharmacology. https://doi.org/10.1038/npp.2015.180

Milad, M.R., Orr, S.P., Lasko, N.B., Chang, Y., Rauch, S.L., Pitman, R.K., 2008. Presence and acquired origin of reduced recall for fear extinction in PTSD: Results of a twin study. J. Psychiatr. Res. 42, 515–520. https://doi.org/10.1016/j.jpsychires.2008.01.017

Milad, M.R., Pitman, R.K., Ellis, C.B., Gold, A.L., Shin, L.M., Lasko, N.B., Zeidan, M.A., Handwerger, K., Orr, S.P., Rauch, S.L., 2009. Neurobiological Basis of Failure to Recall Extinction Memory in Posttraumatic Stress Disorder. Biol. Psychiatry 66, 1075–1082. https://doi.org/10.1016/j.biopsych.2009.06.026

Milad, M.R., Quirk, G.J., 2012. Fear extinction as a model for translational neuroscience: Ten years of progress. Annu. Rev. Psychol. https://doi.org/10.1146/annurev.psych.121208.131631

Milad, M.R., Quirk, G.J., 2002. Neurons in medial prefrontal cortex signal memory for fear extinction. Nature 420, 70–74. https://doi.org/10.1038/nature01138

Norrholm, S.D., Jovanovic, T., Olin, I.W., Sands, L.A., Karapanou, I., Bradley, B., Ressler, K.J., 2011. Fear extinction in traumatized civilians with posttraumatic stress disorder: Relation to symptom severity. Biol. Psychiatry 69, 556–563. https://doi.org/10.1016/j.biopsych.2010.09.013

Overstreet, L.S., Westbrook, G.L., 2003. Synapse Density Regulates Independence at Unitary Inhibitory Synapses.

Piggott, V.M., Bosse, K.E., Lisieski, M.J., Strader, J.A., Stanley, J.A., Conti, A.C., Ghoddoussi, F., Perrine, S.A., 2019. Single-prolonged stress impairs prefrontal cortex control of amygdala and striatum in rats. Front. Behav. Neurosci. 13. https://doi.org/10.3389/fnbeh.2019.00018

Quirk, G.J., Garcia, R., González-Lima, F., 2006. Prefrontal Mechanisms in Extinction of Conditioned Fear. Biol. Psychiatry. https://doi.org/10.1016/j.biopsych.2006.03.010

Quirk, G.J., Likhtik, E., Pelletier, J.G., Paré, D., 2003. Stimulation of medial prefrontal cortex decreases the responsiveness of central amygdala output neurons. J. Neurosci. 23, 8800–7.

Quirk, G.J., Mueller, D., 2008. Neural mechanisms of extinction learning and retrieval. Neuropsychopharmacology 33, 56–72. https://doi.org/10.1038/sj.npp.1301555

Quirk, G.J., Russo, G.K., Barron, J.L., Lebron, K., 2000. The role of ventromedial prefrontal cortex in the recovery of extinguished fear. J. Neurosci. 20, 6225–31.

Ressler, K.J., Rothbaum, B.O., Tannenbaum, L., Anderson, P., Graap, K., Zimand, E., Hodges, L., Davis, M., 2004. Cognitive enhancers as adjuncts to psychotherapy: Use of D-cycloserine in phobic individuals to facilitate extinction of fear. Arch. Gen. Psychiatry 61, 1136–1144. https://doi.org/10.1001/archpsyc.61.11.1136

Rosso, I.M., Weiner, M.R., Crowley, D.J., Silveri, M.M., Rauch, S.L., Jensen, J.E., 2014. Insula and anterior cingulate GABA levels in posttraumatic stress disorder: Preliminary findings using magnetic resonance spectroscopy. Depress. Anxiety 31, 115–123. https://doi.org/10.1002/da.22155

Schofield, C.M., Huguenard, J.R., 2007. GABA affinity shapes IPSCs in thalamic nuclei. J. Neurosci. 27, 7954–7962. https://doi.org/10.1523/JNEUROSCI.0377-07.2007

Schroeder, B.C., Waldegger, S., Fehr, S., Bleich, M., Warth, R., Greger, R., Jentsch, T.J., 2000. A constitutively open potassium channel formed by KCNQ1 and KCNE3. Nature 403, 196–199. https://doi.org/10.1038/35003200

Schür, R.R., Draisma, L.W.R., Wijnen, J.P., Boks, M.P., Koevoets, M.G.J.C., Joëls, M., Klomp, D.W., Kahn, R.S., Vinkers, C.H., 2016. Brain GABA levels across psychiatric disorders: A systematic literature review and meta-analysis of 1H-MRS studies. Hum. Brain Mapp. https://doi.org/10.1002/hbm.23244

Senn, V., Wolff, S.B.E., Herry, C., Grenier, F., Ehrlich, I., Gründemann, J., Fadok, J.P., Müller, C., Letzkus, J.J., Lüthi, A., 2014. Long-range connectivity defines behavioral specificity of amygdala neurons. Neuron 81, 428–437. https://doi.org/10.1016/j.neuron.2013.11.006

Sierra-Mercado, D., Padilla-Coreano, N., Quirk, G.J., 2011. Dissociable roles of prelimbic and infralimbic cortices, ventral hippocampus, and basolateral amygdala in the expression and extinction of conditioned fear. Neuropsychopharmacology 36, 529–538. https://doi.org/10.1038/npp.2010.184

Sotres-Bayon, F., Quirk, G.J., 2010. Prefrontal control of fear: More than just extinction. Curr. Opin. Neurobiol. https://doi.org/10.1016/j.conb.2010.02.005

Souza, R.R., Noble, L.J., McIntyre, C.K., 2017. Using the Single Prolonged Stress Model to Examine the Pathophysiology of PTSD. Front. Pharmacol. 8, 615. https://doi.org/10.3389/fphar.2017.00615

Ting, J.T., Daigle, T.L., Chen, Q., Feng, G., 2014. Acute brain slice methods for adult and aging animals: Application of targeted patch clamp analysis and optogenetics. Methods Mol. Biol. 1183, 221–242. https://doi.org/10.1007/978-1-4939-1096-0_14

Victoria, N.C., Marron Fernandez de Velasco, E., Ostrovskaya, O., Metzger, S., Xia, Z., Kotecki, L., Benneyworth, M.A., Zink, A.N., Martemyanov, K.A., Wickman, K., 2016. G Protein-Gated K+ Channel Ablation in Forebrain Pyramidal Neurons Selectively Impairs Fear Learning. Biol. Psychiatry 80, 796–806. https://doi.org/10.1016/j.biopsych.2015.10.004

Xiong, Z., Zhang, K., Ishima, T., Ren, Q., Chang, L., Chen, J., Hashimoto, K., 2018. Comparison of rapid and long-lasting antidepressant effects of negative modulators of α5-containing GABAA receptors and (R)□ketamine in a chronic social defeat stress model. Pharmacol. Biochem. Behav. 175, 139–145. https://doi.org/10.1016/j.pbb.2018.10.005

Yamamoto, S., Morinobu, S., Fuchikami, M., Kurata, A., Kozuru, T., Yamawaki, S., 2008. Effects of single prolonged stress and D-cycloserine on contextual fear extinction and hippocampal NMDA receptor expression in a rat model of PTSD. Neuropsychopharmacology 33, 2108–2116. https://doi.org/10.1038/sj.npp.1301605

Yuen, E.Y., Liu, W., Karatsoreos, I.N., Feng, J., McEwen, B.S., Yan, Z., 2009. Acute stress enhances glutamatergic transmission in prefrontal cortex and facilitates working memory. Proc. Natl. Acad. Sci. U. S. A. 106, 14075–14079. https://doi.org/10.1073/pnas.0906791106

